# SpaceANOVA: Spatial co-occurrence analysis of cell types in multiplex imaging data using point process and functional ANOVA

**DOI:** 10.1101/2023.07.06.548034

**Authors:** Souvik Seal, Brian Neelon, Peggi Angel, Elizabeth C. O’Quinn, Elizabeth Hill, Thao Vu, Debashis Ghosh, Anand Mehta, Kristin Wallace, Alexander V. Alekseyenko

## Abstract

**Motivation:** Multiplex imaging platforms have enabled the identification of the spatial organization of different types of cells in complex tissue or tumor microenvironment (TME). Exploring the potential variations in the spatial co-occurrence or co-localization of different cell types across distinct tissue or disease classes can provide significant pathological insights, paving the way for intervention strategies. However, the existing methods in this context either rely on stringent statistical assumptions or suffer from a lack of generalizability.

**Results:** We present a highly powerful method to study differential spatial co-occurrence of cell types across multiple tissue or disease groups, based on the theories of the Poisson point process (PPP) and functional analysis of variance (FANOVA). Notably, the method accommodates multiple images per subject and addresses the problem of missing tissue regions, commonly encountered in such a context due to the complex nature of the data-collection procedure. We demonstrate the superior statistical power and robustness of the method in comparison to existing approaches through realistic simulation studies. Furthermore, we apply the method to three real datasets on different diseases collected using different imaging platforms. In particular, one of these datasets reveals novel insights into the spatial characteristics of various types of precursor lesions associated with colorectal cancer.

**Availability:** The associated *R* package can be found here, https://github.com/sealx017/SpaceANOVA.

**Supplementary information:** The supplementary material is attached.

## 1 Introduction

The advent of multiplex tissue imaging technologies (Heinzmann *et al*., 2017; Burlingame *et al*., 2021; Lewis *et al*., 2021; Liu *et al*., 2022; Seal and Ghosh, 2022; Wrobel *et al*., 2023), such as cyclic immunofluorescence (CyCIF) (Lin *et al*., 2015), co-detection by imaging (CODEX) (Goltsev *et al*., 2018), imaging mass cytometry (IMC) (Giesen *et al*., 2014), multiplex immunohistochemistry/immunofluorescence (mIHC/IF) (Tsujikawa *et al*., 2017), and multiplexed ion beam imaging (MIBI) (Angelo *et al*., 2014), has revolutionized our ability to probe the spatial biology of tissues and tumor microenvironments (TMEs) (Binnewies *et al*., 2018) at the single-cell level with unprecedented detail and resolution. These technologies and respective commercial platforms, such as Vectra 3.0, PhenoCycler, Vectra Polaris (all three from Akoya Biosciences), MIBI (Ionpath Inc.), and Visikol Services (BICO), are chosen by researchers based on the pathological context and the specific research questions at hand. The overarching goal is to uncover the intricate mechanisms that dictate cellular and protein interactions and to further examine their association with relevant clinical outcomes. The datasets acquired from such platforms usually share a similar structure. As such, each subject may have one or more non-overlapping two-dimensional images of the tissue or TME. These images are captured at a cellular and nucleus-level resolution, wherein the proteins within the sample have been labeled using antibodies or markers. The number of markers varies based on the platform, e.g., Vectra Polaris images typically have 6-8 markers, whereas MIBI images can have 40 or more markers.

After the initial step of identifying cellular boundaries through single-cell segmentation, different cell types, such as CD4+ T helper cells, CD8+ Cytotoxic T cells, and tumor cells, are detected based on supervised or semi-supervised clustering (Schapiro *et al*., 2017; Seal *et al*., 2022) using the continuous-valued intensity of the respective surface or phenotypic markers (Shipkova and Wieland, 2012). Once the cell types have been identified, their relative abundance and additionally, spatial organization can be studied. In the presence of a clinical phenotype that categorizes subjects into different groups, such as different types of treatments or various sub-types of tissues or tumors, it becomes feasible to explore whether the spatial co-occurrence or interaction of these different cell types varies across the groups. Such an analysis can help to decipher the relationship between cellular interactions and the clinical phenotype, with potential implications for personalized treatment strategies (Färkkilä *et al*., 2020).

Some of the earlier works that involved quantifying spatial co-occurrence of cell types (Schapiro *et al*., 2017; Keren *et al*., 2018; Damond *et al*., 2019; Tsakiroglou *et al*., 2020; Schürch *et al*., 2020), are based on simple yet intuitive measures, such as counting the number of touching cells or, more generally, the number of nearest neighbors of every cell and inspecting the proportion of different cell types. For a formal inference, cell types are randomly switched and empirical *p*-values based on permutation test (Welch, 1990) are considered. Recently, a more rigorous framework based on the homogeneous multitype Poisson point process (PPP) or complete spatial randomness and independence (CSRI) (Diggle, 2013; Baddeley *et al*., 2015) has been considered by different groups of researchers (Bull *et al*., 2020; Vipond *et al*., 2021; Wilson *et al*., 2022; Canete *et al*., 2022; Vu *et al*., 2022). In essence, these methods utilize various spatial summary functions, such as Ripley’s K (Ripley, 1976), L, g, and mark connection function (mcf), which are all based on the same set of assumptions and have slightly varying interpretations. As an example, for a pair of cell types (*m, m*′), the bivariate K-function (*K*_*mm′*_ (*r*)) calculates the ratio of the observed number of cells to the expected number of cells of type *m*′ within a distance *r* of a typical cell of type *m*, relative to a hypothetical completely random distribution of cell types. It is worth highlighting that variations of these summary functions have been historically used in diverse fields like ecology (Legendre and Fortin, 1989), epidemiology (Gatrell *et al*., 1996), astronomy (Kerscher, 2000), and crime research (Anselin *et al*., 2000). In the context of general spatial pathology datasets, many recent works (Osher *et al*., 2023; Dayao *et al*., 2023) are using innovative modeling frameworks based on PPP to answer different spatial questions.

With the assumptions of homogeneous multitype PPP or CSRI for studying cellular co-occurrence, most of the relevant works simplify the analysis of summary functions by reducing them to simple univariate statistics. For example, Wilson *et al*. (2022) picks a specific value of the radius *r* based on clinical relevance and considers *K*_*mm′*_ (*r*) for different pairs of cell types. In the *R*-package called SpicyR, Canete *et al*. (2022) considers a numerical approximation of the integrated value of the L function, by summing up the function for a few chosen values of *r*. Such a simplification of the summary functions, however, discards granular information and thus, might be sub-optimal in practical scenarios, as demonstrated in our simulation studies. A key improvement was proposed by Vu *et al*. (2022) who incorporates the mark connection function (mcf) over a range of *r*, to the additive functional Cox model (Cui *et al*., 2021), as a functional covariate, to study association with time-to-event outcomes. However, unlike SpicyR, Vu *et al*. (2022)’s approach can not be readily used to study differential spatial co-occurrence across clinical groups, which is the focus of our manuscript and also, does not accommodate more than one image per subject. Additionally, the multiplexed images may have holes in them, i.e., some portions of the tissue can be missing, largely due to the intricacies related to data collection procedures. In such a scenario, directly employing assumptions of multitype PPP or CSRI can lead to spurious and artifactual associations, as acknowledged by Wilson *et al*. (2022).

Practically addressing the aforementioned limitations, we propose a new method termed SpaceANOVA. For every pair of cell types (*m, m*′), we consider the pair correlation function or g-function (*g*_*mm′*_ (*r*)) that stands for the probability of observing a cell of type *m*′ at an exact distance of *r* from a usual point of type *m*, divided by the corresponding probability under a homogenous multitype PPP. Then, we compare the functional values: *g*_*mm′*_ (*r*) over a range of *r*, across subjects in different groups using a functional analysis of variance or FANOVA framework (Zhang, 2013; Smaga and Zhang, 2019). The rationale behind focusing on the g-function is its ease of interpretation in comparison to the K or L functions, which are “cumulative” in nature (Baddeley *et al*., 2015). To mitigate the bias due to holes or missing areas in the tissue, we adjust the standard g-function by a permutation-based envelope (Baddeley *et al*., 2014), a variation of which is used in Wilson *et al*. (2022). To handle multiple images per subject, we evaluate two options, firstly, a one-way FANOVA model with the simple or weighted average of the g-functions corresponding to the images per subject, and secondly, a modified two-way FANOVA model considering every g-function individually. We demonstrate SpaceANOVA’s robustness and superior power as compared to the popular method named SpicyR, in realistic simulation scenarios, one of which focuses on simulating images with missing regions or holes and highlights the need to account for this problem with great care. We employ SpaceANOVA to spatially differentiate between distinct types of colorectal polyps (Wallace *et al*., 2021) by analyzing an mIF dataset collected using the Vectra Polaris platform (Akoya Biosciences) at the Medical University of South Carolina (MUSC). In addition to that, we analyze two publicly available datasets: an IMC dataset on type 1 diabetes (Damond *et al*., 2019) and a colorectal cancer dataset collected using the CODEX platform (Akoya Biosciences) (Schürch *et al*., 2020). SpaceANOVA is available as an *R*-package at this link, https://github.com/sealx017/SpaceANOVA.

## 2 Materials and Methods

### 2.1 Assumptions

Suppose there are *M* distinct cell types and *G* clinical groups with group *g* having *s*_*g*_ subjects for *g* = 1, …, *G*. Denote the number of images of a subject *i* from the group *g* to be *n*_*gi*_ and the total numbers of images to be 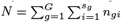. Each image may have an arbitrary number of cells.

Under the hypothesis of no spatial co-occurrence between any pair of cell types, we make two crucial assumptions. First, we assume that the cells of any type *m* constitute a homogeneous Poisson point process (PPP) or complete spatial randomness (CSR), with a constant intensity of *λ*_*m*_ in every image (omitting image or subject indices for notational simplicity). In simple terms, this assumption of CSR tells us that for any cell type *m*, the corresponding cells have no preference for any spatial location, and information about the cells in one region of the tissue has no influence on other regions. Further, we assume that the cells of different types are independent. With these two assumptions considered collectively, the entire setup constitutes a homogeneous multitype PPP or complete spatial randomness and independence (CSRI) (Baddeley *et al*., 2015).

### 2.2 Quantification of spatial co-occurrence

For any multitype point process (not necessarily, homogeneous or Poisson), the spatial correlation or co-occurrence between two types (*m, m*′), can be studied using the bivariate version of Ripley’s *K* function (Ripley, 1976) as defined below,

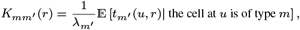

where 𝔼 denotes the expectation operator and *t*_*m′*_ (*u, r*) denotes the number of cells of type *m*′ lying within a circle of radius *r* from a location *u* that has a cell of type *m*. Similarly, the bivariate pair correlation or g-function can be defined as, 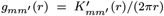, where 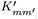 denotes the first derivative of *K*_*mm′*_. It should be highlighted that *g*_*mm′*_ (*r*), unlike *K*_*mm′*_ (*r*), is not cumulative in nature, i.e., *g*_*mm′*_ (*r*) contains contributions only from interpoint distances exactly equal to *r*, while K-function detects interaction that occurs equally at all distances up to a certain maximum distance *r*. Under our assumptions of homogeneous multitype PPP or CSRI, it can be shown that *K*_*mm′*_ (*r*) = *πr*^2^, and *g*_*mm′*_ (*r*) = 1, for all *r* ∈ [0, *R*], and for any *m, m*′ ∈ {1, 2, …, *M* }.

In a real dataset, the K-function of each image can be estimated as,

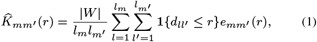

where *l*_*m*_ and *l*_*m′*_ denote the number of cells of types *m* and *m*′ in the image, respectively. *d*_*ll′*_ denotes the distance between the *l*-th cell of type *m* and *l*′-th cell of type *m*′. |*W* | denotes the image area and *e*_*mm′*_ (*r*) is an edge correction factor accounting for irregular image boundary. The g-function, 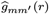 can be estimated using numerical differentiation of 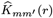. When the estimated summary functions vary significantly from their expected values under CSRI, one can conclude that there is evidence of spatial correlation or co-occurrence between cell types.

In the next section, we propose two modeling frameworks based on functional analysis of variance or FANOVA to compare the estimated g-functions across groups. It is worth mentioning that investigating K or g-functions to identify sub-clusters or sub-populations using different approaches is historically popular in ecological research, e.g., Illian *et al*. (2006) performs functional principal component analysis (FPCA) of g-functions of multiple plant species to group them on the basis of their scores on the principal components.

### 2.3 Functional analysis of variance (FANOVA)

In this section, for each pair of cell types, we will test if the estimated g-functions of the images from subjects across *G* groups exhibit any variation or differential pattern.

Let *Y*_*gij*_ (*r*) denote the value of the estimated g-function at a radius *r* of image *j* of a subject *i* belonging to group *g*: 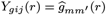 (the group index *g* is not to be confused with the g-function). Each pair of cell types (*m, m*′) is analyzed individually, and thus, the notations *m* and *m*′ are henceforth not used.

#### 2.3.1 Univariate approach

Consider the simplest case where every subject has a single image, i.e., *n*_*gi*_ = 1 for all *i* and *g* (and thus, 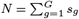). Next, we consider the functional analog of a standard one-way ANOVA model, also known as FANOVA (Zhang, 2013) in the following way (dropping the image index *j*),

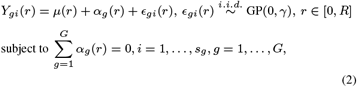

where *μ*(*r*), *α*_*g*_(*r*), *ϵ*_*gi*_(*r*), denote the common, group-specific, and subject-specific mean functions, respectively. GP(0, *γ*) denotes a Gaussian process with zero mean function and covariance function *γ*(*r*_1_, *r*_2_) for *r*_1_, *r*_2_ ∈ [0, *R*] (Banerjee *et al*., 2008). Additional assumptions are provided in the supplementary material.

The null hypothesis (*H*_0_) of an equal level of spatial co-occurrence at any value of the radius *r* across *G* groups can be formulated as,

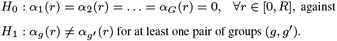

The total sum of squares (SS_*T*_) at a particular *r* ∈ [0, *R*] can be partitioned into two components as,

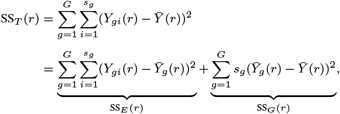

where SS_*E*_ (*r*) and SS_*G*_(*r*) stand for within-groups and between-groups variability respectively. To test *H*_0_, we consider the global pointwise *F* - type test statistic (GPF) (Smaga and Zhang, 2019) given by the formula,

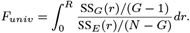

The null distribution of *F*_*univ*_ can be approximated by a 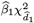 distribution, where 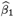 and 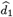 are suitably estimated from the data and their forms and derivations are provided in the supplementary material.

Note that this setup can be readily used when we have multiple images per subject (*n*_*gi*_ ≥ 1) by considering a simple mean of the functions of different images corresponding to each subject, i.e., replacing *Y*_*gi*_(*r*) by 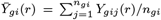 above. The crucial assumption that we would be making with such a simplification, is that the images from a particular subject are different realizations of the same homogeneous multitype PPP. This assumption might not be implausible, especially if the images correspond to different regions of the same tissue or TME, as in the majority of the realistic scenarios. In the results section, we refer to this approach as “SpaceANOVA Univ.”.

#### 2.3.2 Multivariate approach

In addition to the mean-based approach, when subjects have more than one image, a more sophisticated way would be to extend the one-way FANOVA model from Equation (2) in the following way,

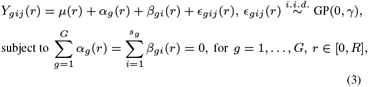

where *μ*(*r*), *α*_*g*_(*r*), *β*_*gi*_(*r*) and *ϵ*_*gij*_ (*r*), denote the common, group-specific, subject-specific, and image-specific mean functions, respectively.

The total sum of squares can be partitioned as,

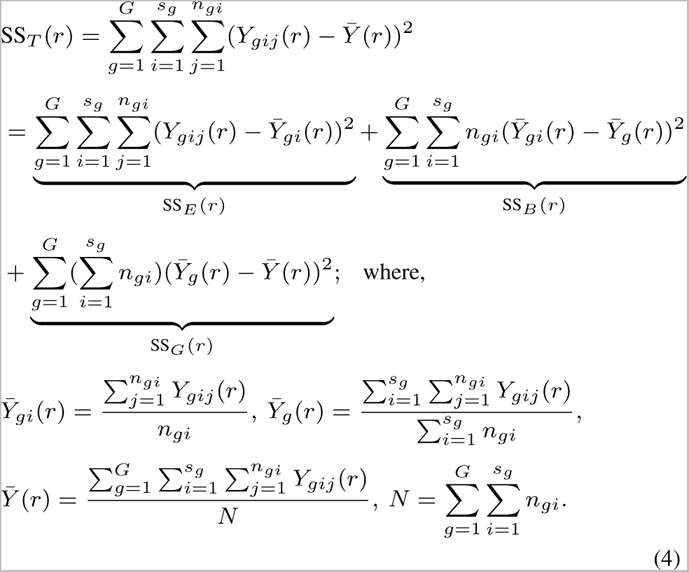

A test statistic analogous to *F*_*univ*_ can be formulated as,

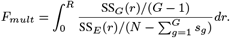

The null distribution of *F*_*mult*_ can be approximated by a 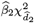 distribution, where 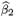 and 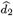 are suitably estimated from the data and their forms and additional information are provided in the supplementary material. In the results section, we refer to this approach as “SpaceANOVA Mult.”.

The integrals in both *F*_*univ*_ and *F*_*mult*_ are approximated using a numerical integration where *r* is varied across a fixed grid of values between 0 and a pre-selected large value of *R*.

### 2.4 Adjustment to account for missing tissue regions

During the imaging data-collection procedure, some areas of the tissue or TME can become folded or torn because of slicing. Thus, the resulting images can have holes or blank regions in them where no cells are observed. For such images, the assumption of homogeneous multitype PPP can be too restrictive since the holes introduce an additional degree of sparseness or “in-homogeneity”. It can further lead to spatial patterns of cell types that are mere artifacts (see Simulation section).

As alluded, one possible solution can be to relax the assumption of homogeneity to consider inhomogeneous multitype PPP instead. It would assume that the intensity of a cell type *m* is not constant and varies w.r.t. to any location *z* of an image *W*, i.e., *λ*_*m*_(*z*) ≠ *λ*_*m*_ for *z* ∈ *W*. However, the estimation of *λ*_*m*_(*z*) is a difficult task, one that can further bias the estimation of the K and g-functions down the line (Baddeley *et al*., 2015). Thus, we allow the assumption of homogeneity and consider a permutation envelope-based adjustment instead. To elaborate, we randomly switch the cell type labels in each image a fixed number of times (say, *P* = 50) and compute 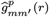 every time. Then, we adjust the original g-function 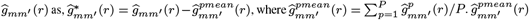 can be interpreted as the “true” expected value of 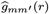 under the null hypothesis of no spatial co-occurrence instead of the value 1, which is expected only when an image does not suffer from an additional source of heterogeneity. Thus, the adjusted g-function would represent the spatial correlation that is not caused spuriously by the non-uniform or inhomogeneous structure of the image but is due to the process’s deviation from CSRI. A variation of such an adjustment has been used in Wilson *et al*. (2022). Originally, Baddeley *et al*. (2014) introduced the idea of a permutation-based envelope for testing departure from CSR in a simpler context.

## 3 Real Data Analysis

We applied permutation-adjusted versions of SpaceANOVA Univ. and SpaceANOVA Mult. to the following real datasets,

1. A multiplex immunofluorescence (mIF) dataset on colorectal adenoma obtained using the Vectra Polaris platform (Akoya Biosciences) at the MUSC.
2. An imaging mass cytometry (IMC) dataset on type 1 diabetes mellitus (T1DM) (Damond *et al*., 2019).
3. An mIF dataset on colorectal cancer (CRC) collected using the CODEX platform (Akoya Biosciences) (Schürch *et al*., 2020).

To maintain the manuscript’s brevity, the figures corresponding to the analysis of the T1DM dataset and the full analysis of the CRC dataset are provided in the supplementary material. The radius *r* was varied between [0, 100] with a step of size 1, i.e., *r* = 0, 1, 2, …, 100 in the first two analyses.

### 3.1 Application to colorectal polyps dataset

In the colorectal polyps dataset, there are 19 subjects each with a polyp of type either sessile serrated adenoma (SSA) or tubular/villous adenoma (TA/VA) (Nagtegaal *et al*., 2020). Every subject has varying numbers of images (between 15-380). There are six markers: CD4, CD8, CK, Foxp3, RORgt, and Tbet, based on which different cell types, such as Th (CD4+), Tc (CD8+), Treg (CD4+Foxp3+), and Th17 (CD4+RORgt+), are identified. We applied adjusted SpaceANOVA Univ. and SpaceANOVA Mult. to understand how the pair-wise spatial co-occurrence of these four immune cell types: Th, Tc, Treg, and Th17, differ between the two groups of polyps. Refer to the supplementary material for details about the data collection procedures.

From Figure 2, notice that the cell type pairs such as (Tc, Tc), (Th, Tc), and (Treg, Th), were detected by both the methods (more prominently by SpaceANOVA Mult.) to exhibit differential spatial co-occurrence between the two groups at a level *α* = 0.05. In Figure 3, for pairs: (Treg, Th) and (Th, Tc), the mean g-function (over images) of every subject *i* from group 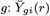 (from Equation 4) and the overall group means: 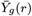 (from Equation 4) are displayed. For (Treg, Th), the g-functions in both the groups seemingly peaked at around the radius *r* = 10, meaning that surrounding any Treg cell, the highest probability of finding another Th cell is at a distance of 10. The group mean function of TA/VA appeared to be slightly higher than that of SSA for all *r* ∈ [0, 100], implying that the Treg cells were spatially closer to Th cells in TA/VA subjects as opposed to SSA subjects. For (Th, Tc), the g-functions were mostly negative, especially for smaller values of *r*, implying that these two cell types avoided each other or showed decreased spatial co-occurrence. Furthermore, comparing the group mean function of SSA to that of TA/VA, the former exhibited more negative values. It suggested that Th and Tc cells in SSA subjects had a stronger tendency of decreased spatial co-occurrence. In Figure 3C and 3D, the spatial organization of cells in two representative images from two subjects of the two groups is shown. The figures supported the previous conclusions. Specifically, the images from the TA/VA subject showed increased co-occurrence of Treg cells, while in the images from the SSA subject, Th and Tc cells appeared to be avoiding or located further away from each other.

**Fig. 1.**
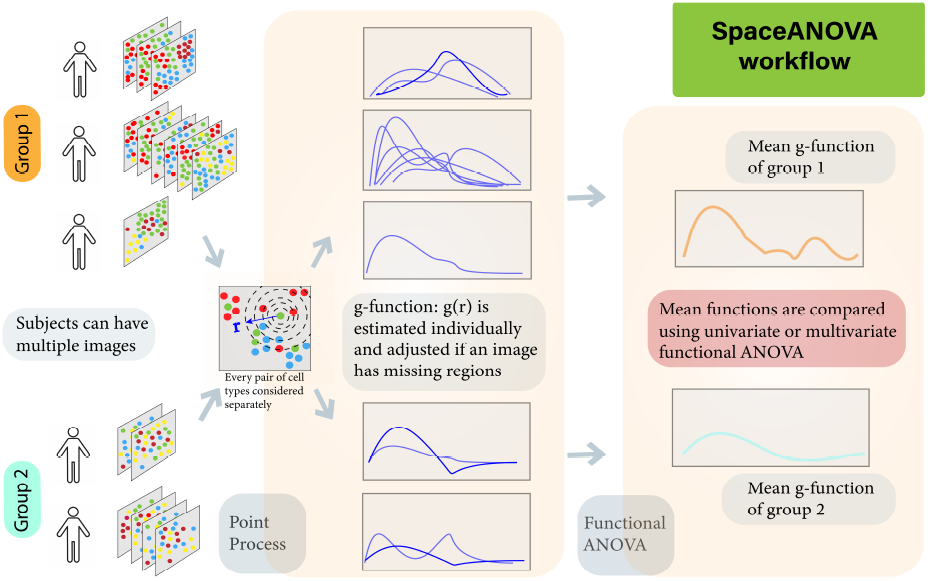
The workflow of the proposed method. Assumptions of spatial point process are used to summarise the level of spatial co-occurrence of pairs of cell types in each image of the subjects from different groups in the form of g-function and compared either using a univariate or multivariate functional ANOVA approach. Although only two groups are depicted in the figure, the method can handle any number of groups, i.e., *G* ≥ 2.

**Fig. 2.**
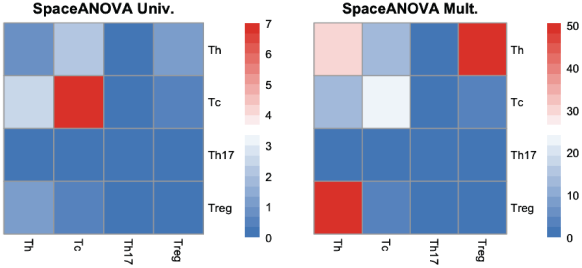
Heatmap of - log_10_ (*p*)-values of the two methods for every pair of the four immune cell types: Th, Tc, Treg, and Th17.

**Fig. 3.**
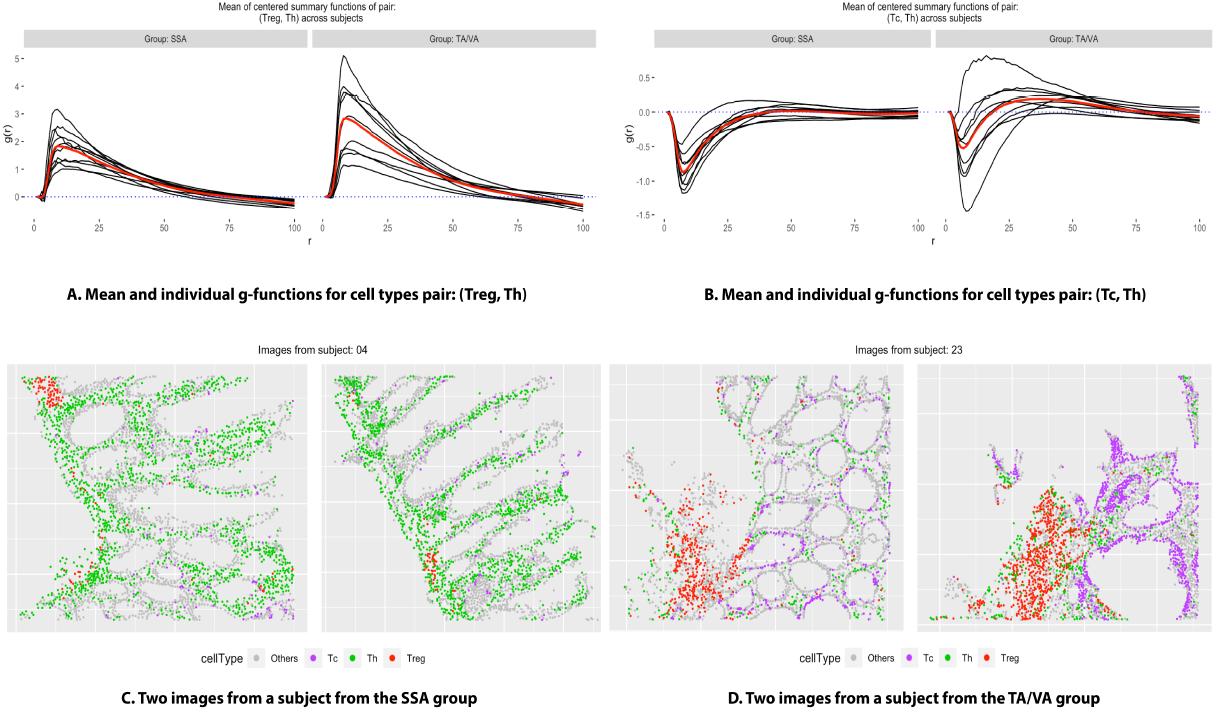
Figure (A) shows the individual g-function (averaged over images) of every subject in two groups (in the color black) and also the group mean function (in the color red), for the pair of cell types: (Treg, Treg). Figure (B) shows a similar plot for the pair: (Tc, Th). Figures (C) and (D) show the cellular organization of representative images from two subjects from the groups SSA and TA/VA, respectively, where different colors correspond to different cell types.

The spatial relationship of T-cells in preinvasive colorectal lesions of different histologic types has not previously been investigated. The Treg-Th co-occurrence pattern we observed among conventional adenomas suggests a persistent immune suppressive environment. Th cells direct the adaptive immune response within the TME, determining an anti-tumor cytotoxic or anti-inflammatory regulatory response, potentially providing a growth advantage to the developing lesion (Rozek *et al*., 2016). Additional information on the CD4+Foxp3+ lineage may clarify the nature of Tregs in promoting or inhibiting tumor growth in conventional lesions (Maoz *et al*., 2019). For example, a CD4+Th17+Foxp3+ population is highly immune suppressive (Galon *et al*., 2006) and tumor-promoting whereas other CD4+Foxp3+ populations are associated with better prognosis (De Smedt *et al*., 2015). The inverse association between Tc and Th populations in SSA likely reflects differences in types and functions of lymphoid clustering (Rozek *et al*., 2016). For example, the Crohn’s-like lymphoid reaction occurs peritumorally, is often densely populated with Th cells, and as it matures, assists with lymphocyte recruitment to the tumor bed. Clusters of tumor-infiltrating lymphocytes (TILs), on the other hand, emerge within the tumor nest and are typically comprised of Tc populations designed to kill neoplastic cells Galon *et al*. (2006). Interestingly, these lymphoid clustering patterns are also observed within microsatellite stable (MSS) and microsatellite instable (MSI+) colorectal cancer (CRC) (De Smedt *et al*., 2015; Chang *et al*., 2018), the putative parent lesions to SSA, suggesting a shared immunologic clustering lineage. In future studies, it will be important to co-localize immune clustering to pathologically annotated regions (e.g. aggregates, dysplasia) to better inform the spatial immune clusters.

### 3.2 Application to the type 1 diabetes dataset

In the type 1 diabetes mellitus (T1DM) dataset (Damond *et al*., 2019), there are 8 subjects, four of which belong to the non-diabetic group and the rest belong to the onset group. Each subject has varying numbers of images (between 64-81) and cells of sixteen different types of which we focused on studying the pairwise spatial co-occurrence of alpha, beta, delta, Th, Tc, neutrophil, and macrophage cells. In Figure S1 of the supplementary material, the heatmaps of - log_10_(*p*)-values for different pairs of cell types are displayed. At a level *α* = 0.05, SpaceANOVA Univ. only detected two pairs: (beta, beta) and (delta, delta) to exhibit differential spatial co-occurrence, while SpaceANOVA Mult. detected several more such as (alpha, delta), (beta, delta), and (Tc, beta). In Figure S2 of the supplementary material, the g-functions corresponding to the pairs (alpha, delta) and (Tc, beta) are displayed. For (alpha, delta), we noted that the g-functions of both groups were mostly positive, particularly for smaller values of *r*, indicating an increased or positive spatial co-occurrence. Additionally, the g-functions of the onset group attained slightly higher values than the non-diabetic group implying that the cell types showed higher spatial co-occurrence in the former. The finding agreed with similar works in this field such as Gao *et al*. (2021)’s study of delta cells that revealed delta cells’ significant interaction with other endocrine cell types, alpha and beta in the context of T1DM. For (Tc, beta), we noted that the g-functions of both groups were negative for smaller values of the radius *r* and became close to 0 and slightly higher as *r* increased. It indicated avoidance or decreased spatial co-occurrence of Tc and beta cells, particularly in the non-diabetic group. This result was in accordance with Damond *et al*. (2019)’s significant discovery using the same dataset that Tc cells interact less with the beta cells in normal subjects but start to infiltrate the beta cell-rich areas more as T1DM progresses.

## 4 Simulation

We evaluated the performance of both the univariate and multivariate versions of our proposed method: SpaceANOVA Univ. and SpaceANOVA Multi. in different simulation setups. For comparison, we included the two tests provided in the popular *R*-package SpicyR (Canete *et al*., 2022): one based on a linear model and the other based on a linear mixed model, called SpicyR-LM and SpicyR-Mixed, respectively. Note that, the g-functions were not adjusted for missing tissue regions in the first two simulations, while in the third simulation setup, we added the adjusted versions: SpaceANOVA Univ. (Adjusted) and SpaceANOVA Multi. (Adjusted) in the comparison. In all three simulations, the radius *r* was varied between [0, 200] with a step of size 1, i.e., *r* = 0, 1, 2, …, 200.

### 4.1 Simulation based on mixed Poisson process

In this simulation setup, we considered a variation of the framework described in Canete *et al*. (2022), which can be loosely interpreted as a mixed Poisson point process (Baddeley *et al*., 2015). There were two groups of subjects, each with *N/*2 subjects where two values of *N* : 50 and 100 were considered. In multiplex imaging datasets, usually, a large tissue or organ is divided into smaller non-overlapping parts for easier and more feasible profiling, which results in multiple images per subject. Aligning with that principle, we assumed that each subject had three images of equal area, coming from a “super”-image of size 1000 × 1000 sq. units. There were two cell types A, B with each type having 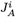 and 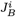 numbers of cells in a subject *i*, where 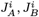 were randomly-drawn integers from from [200, 400]. To generate the super-image, the cells of type A were simulated first following a homogeneous PPP with an intensity equal to the ratio of 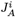 and the image area (= 1000^2^). Next, the density (or, the space-varying intensity) of cell type A was calculated using a disc kernel, where the size of the disc for each subject was either Sigma or (Sigma + Sigma-difference) depending on which group that subject belonged to. The cells of type B were then generated using an inhomogeneous PPP with the estimated density of cell type A. Different values of Sigma and Sigma-difference were considered. Note that a higher value of sigma implies lower co-occurrence or co-localization. Further details and associated plots about the data-generating process are provided in the supplementary material.

From Figure 4A, we noticed that when the spatial co-occurrence was more prominent, i.e., Sigma was smaller (≤ 40), all the methods performed quite close to each other. However, as Sigma increased, the higher power of both SpcaeANOVA Univ. and SpcaeANOVA Mult. became apparent compared to SpicyR-LM and SpicyR-mixed. It could be partially explained by the fact that SpicyR compared the integrated value of the cumulative summary function L between the two groups. In this particular simulation, the L-functions corresponding to the two groups became very close to each other as Sigma increased, while the g-functions kept exhibiting clear differences (refer to the supplementary Figures S4 and S5). Thus, our methods continually achieved superior power. As the number of patients increased from 50 to 100, expectedly, all the methods achieved higher power.

**Fig. 4.**
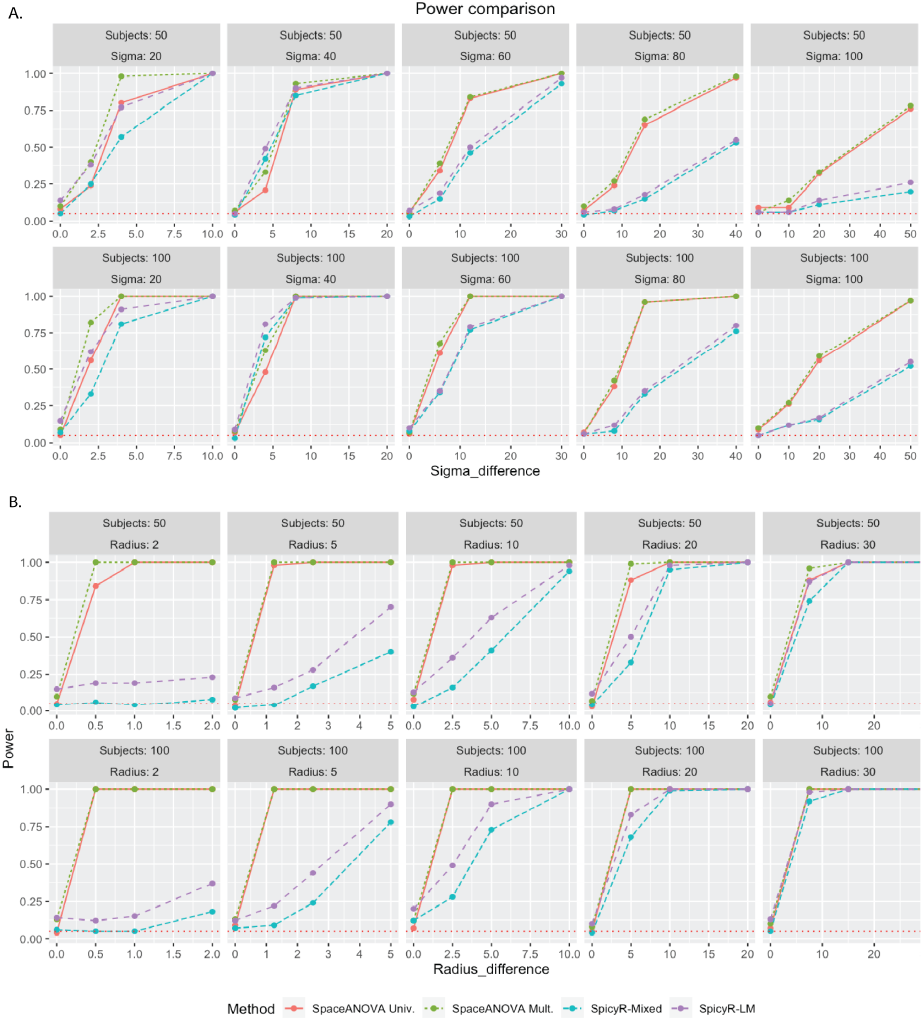
Figures (A) and (B) display the power comparison of the methods in simulation setups based on mixed Poisson process and Netman-Scott cluster process described in the sections 4.1 and 4.2, respectively. The horizontal red-dotted line in every subfigure corresponds to the level *α* = 0.05.

### 4.2 Simulation based on Neyman-Scott cluster process

In this simulation, we followed a similar approach as the previous section altering only the last few steps where the cells of two types were generated. This time, the cells were generated using a multitype Neyman-Scott cluster process (Baddeley *et al*., 2015) with two types A and B. In each super-image, first, a set of “parent” cells was drawn from a homogeneous PPP (without any type) with an intensity equal to the ratio of *J* and the image area (= 1000^2^), where *J* was an integer between [200, 400]. Surrounding each of those parent cells, a cluster of “offspring” cells was generated. Each cluster contained a Poisson number of random cells (with a mean of 3) uniformly distributed either in a disc of radius - “Radius” or a disc of radius -”Radius + Radius-difference” depending on which group the subject belonged to. The parent cells were deleted, and the cells of a cluster were independently marked as type A or B with equal probability. The parameters: Radius and Radius-difference, were varied from low to high. Note that, a larger value of Radius implied higher spatial co-occurrence of the cell types.

From Figure 4B, we noticed that when Radius was smaller (≤ 10), i.e., the degree of spatial co-occurrence was higher, SpcaeANOVA Univ. and SpcaeANOVA Mult. significantly outperformed SpicyR-LM and SpicyR-mixed, while for larger values of Radius, the latter two nearly caught up. This observation was similar to the previous simulation setup and could be explained using a similar argument as well (refer to the supplementary Figures S4 and S5). One additional point to be highlighted is that, in most of the cases, SpicyR-LM produced inflated type-1 errors. SpaceANOVA Mult. displayed slight inflation in the type-1 error in a few cases, while SpaceANOVA Univ. and SpicyR-mixed were nearly perfect in this regard.

### 4.3 Simulation in the presence of missing tissue regions

In this simulation, we followed exactly the same setup described in Section 4.1 with an additional condition that the super-images of subjects from group 1 would have five randomly created circular holes in them. This time, we also considered the adjusted versions of our tests: SpaceANOVA Univ. (Adjusted) and SpaceANOVA Mult. (Adjusted) following the steps outlined in Section 2.4 with *P* = 50.

From Figure 5, notice that all of the previous four methods produced extremely inflated type 1 errors (close to 1), while the adjusted methods produced controlled type 1 error and were adequately powerful in all the cases. It demonstrated the robustness of the proposed permutation envelope-based adjustment and highlighted the necessity of taking into account tissue in-homogeneity under the presence of holes or missing regions in the tissue or TME.

**Fig. 5.**
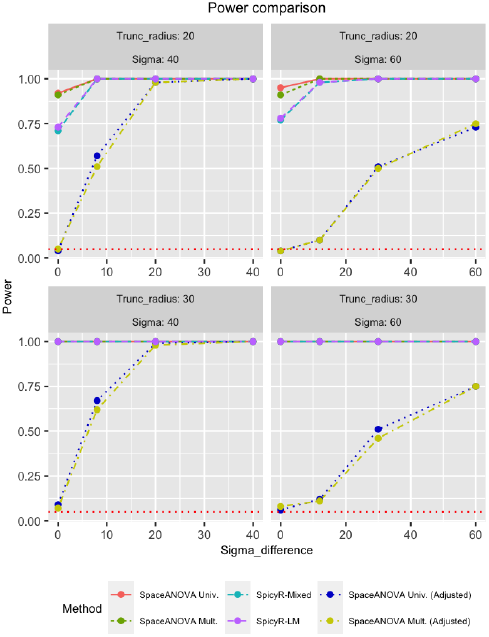
The figure displays the power comparison of the methods in the simulation setup described in the section 4.3 involving missing tissue regions or holes. The horizontal red-dotted line in every subfigure corresponds to the level *α* = 0.05.

## 5 Discussion

In multiplex imaging datasets, studying the spatial co-occurrence or co-localization of different cell types is a crucial task that can provide a key understanding of the complex spatial biology of tissue or TME. When there are multiple groups of subjects based on a pathological phenotype, e.g., case/control or different types of tissues, it becomes further possible to explore if the level of spatial co-occurrence of different pairs of cell types varies across the groups. Such a study of differential spatial co-occurrence could be particularly important to comprehend the disease pathology by revealing the structural and functional differences across the groups.

In this context, we propose a powerful method named SpaceANOVA based on the concepts of homogeneous multitype Poisson point process (PPP) and functional analysis of variance (FANOVA). SpaceANOVA has two variations, namely SpaceANOVA Univ. and SpaceANOVA Mult., which compute a pairwise spatial summary function of cell types in images of every subject individually and then compare the functions across groups using univariate and multivariate versions of FANOVA, respectively. These tests are more general than existing approaches which rely on oversimplifying different spatial summary functions and thereby lose power as demonstrated in our simulations. In addition, SpaceANOVA addresses the issue of images having holes or missing areas in them due to the intricacies of the data-collection procedure, using a permutation envelope-based adjustment. In a practical simulation setup, we show that not accounting for this particular issue might heavily bias all the methods.

We have applied the proposed method to three real datasets, 1) a multiplex immunofluorescence (mIF) dataset on colorectal adenomas and sessile serrated lesions, 2) an IMC dataset on type 1 diabetes mellitus (T1DM), and 3) an IMC dataset on breast cancer. Studying the spatial and functional properties of colorectal neoplasia is a relatively new area of research (Lin *et al*., 2023). This particular mIF dataset has been collected at the Medical University of South Carolina (MUSC) and is currently ongoing thorough statistical dissection that could potentially lead to seminal discoveries. In this dataset, subjects either have a sessile serrated adenoma (SSA) or tubular/villous adenoma (TA/VA) polyp. From a statistical point of view, the dataset is both intriguing and challenging because a) every subject has varying numbers of images (between, 15-380), and b) the images are heavily plagued with the problem of missing tissue regions. We compared the spatial co-occurrence of immune cell types: Th, Tc, Treg, and Th17 between two groups of polyps, SSA and TA/VA. We found several interesting pairs of cell types to exhibit varying levels of spatial co-occurrence between these two groups. For example, Treg cells were found to be surrounded by Th cells in both groups, especially in the TA/VA polyps. On the other hand, Th and Tc cells exhibited avoidance or decreased spatial co-occurrence in both groups, more so in the SSA polyps. We further explored the pathological relevance of these novel findings and established concordance with existing literature. In summary, the real data analyses demonstrate our method’s capability of discovering unique spatial associations between cell types in complex multiplex imaging datasets.

One limitation of the proposed multivariate approach is that it assumes the g-functions corresponding to different images of the same subject to be independent as they only share a common mean and no additional variance components to account for higher-order dependencies. To address this issue, instead of the two-way FANOVA model, we have tried a penalized function-on-function regression (pffr) model (Ivanescu *et al*., 2015) with the group label as a factor variable using a popular *R* package for functional data analysis called Refund (Goldsmith *et al*., 2012). However, the model did not perform as expected because the package is admittedly not well optimized for handling a factor variable. We would like to pursue such a modeling framework with further attention in the future. Also, we have not explicitly discussed adjusting for covariates like age, sex, race, and other pathologic characteristics in our proposed models. One simplistic approach would be to add the covariates directly to the mean, following the form of a simple function-on-scaler regression model (Ramsay and Silverman, 2005). However, when some of the covariates are factor variables, there might be challenges with the interpretation which we will investigate in the upcoming works.

## Software

Software in the form of an *R* package named SpaceANOVA, together with simulation and real data analysis codes are available on GitHub at this link, https://github.com/sealx017/SpaceANOVA. The publicly available datasets are provided as .rda files in the GitHub repository or can be accessed using their original manuscript links.

## Acknowledgments

S.S., B.N., and E.H. were supported in part by the Biostatistics Shared Resource, Hollings Cancer Center, Medical University of South Carolina (P30 CA138313). S.S. and K.W. were supported by NCI R01 CA226086. E.C.O. was supported in part by the Translational Science Shared Resource, Hollings Cancer Center, Medical University of South Carolina (P30 CA138313). A.A. was supported by Quantitative Methods Shared Resource (QMSR) of the South Carolina Cancer Disparities Center (SC CADRE) NCI U54 CA210962 and NLM R01 LM012517. We acknowledge Dr. David N. Lewin’s help with the collection of the colorectal polyps dataset.

## References

Angelo, M. et al. (2014). Multiplexed ion beam imaging of human breast tumors. Nature Medicine, 20(4), 436–442.

Anselin, L., Cohen, J., Cook, D., Gorr, W., and Tita, G. (2000). Spatial analyses of crime. Criminal justice, 4(2), 213–262.

Baddeley, A., Diggle, P. J., Hardegen, A., Lawrence, T., Milne, R. K., and Nair, G. (2014). On tests of spatial pattern based on simulation envelopes. Ecological Monographs, 84(3), 477–489.

Baddeley, A., Rubak, E., and Turner, R. (2015). Spatial point patterns: methodology and applications with R. CRC press.

Banerjee, S., Gelfand, A. E., Finley, A. O., and Sang, H. (2008). Gaussian predictive process models for large spatial data sets. Journal of the Royal Statistical Society: Series B (Statistical Methodology), 70(4), 825–848.

Binnewies, M. et al. (2018). Understanding the tumor immune microenvironment (time) for effective therapy. Nature Medicine, 24(5), 541–550.

Bull, J. A., Macklin, P. S., Quaiser, T., Braun, F., Waters, S. L., Pugh, C. W., and Byrne, H. M. (2020). Combining multiple spatial statistics enhances the description of immune cell localisation within tumours. Scientific reports, 10(1), 18624.

Burlingame, E. A., Eng, J., Thibault, G., Chin, K., Gray, J. W., and Chang, Y. H. (2021). Toward reproducible, scalable, and robust data analysis across multiplex tissue imaging platforms. Cell reports methods, 1(4), 100053.

Canete, N. P., Iyengar, S. S., Ormerod, J. T., Baharlou, H., Harman, A. N., and Patrick, E. (2022). spicyr: Spatial analysis of in situ cytometry data in r. Bioinformatics, 38(11), 3099–3105.

Chang, K., Willis, J., Reumers, J., Taggart, M., San Lucas, F., Thirumurthi, S., Kanth, P., Delker, D., Hagedorn, C., Lynch, P., et al. (2018). Colorectal premalignancy is associated with consensus molecular subtypes 1 and 2. Annals of Oncology, 29(10), 2061–2067.

Cui, E., Crainiceanu, C. M., and Leroux, A. (2021). Additive functional cox model. Journal of Computational and Graphical Statistics, 30(3), 780–793.

Damond, N., Engler, S., Zanotelli, V. R., Schapiro, D., Wasserfall, C. H., Kusmartseva, I., Nick, H. S., Thorel, F., Herrera, P. L., Atkinson, M. A., et al. (2019). A map of human type 1 diabetes progression by imaging mass cytometry. Cell metabolism, 29(3), 755–768.

Dayao, M. T. et al. (2023). Deriving spatial features from in situ proteomics imaging to enhance cancer survival analysis. Bioinformatics, 39(Supplement_1), i140–i148.

De Smedt, L. et al. (2015). Microsatellite instable vs stable colon carcinomas: analysis of tumour heterogeneity, inflammation and angiogenesis. British journal of cancer, 113(3), 500–509.

Diggle, P. J. (2013). i>Statistical analysis of spatial and spatio-temporal point patterns. CRC press.

Färkkilä, A., Gulhan, D. C., Casado, J., Jacobson, C. A., Nguyen, H., Kochupurakkal, B., Maliga, Z., Yapp, C., Chen, Y.-A., Schapiro, D., et al. (2020). Immunogenomic profiling determines responses to combined parp and pd-1 inhibition in ovarian cancer. Nature communications, 11(1), 1459.

Galon, J. et al. (2006). Type, density, and location of immune cells within human colorectal tumors predict clinical outcome. Science, 313(5795), 1960–1964.

Gao, R., Yang, T., and Zhang, Q. (2021). d-cells: The neighborhood watch in the islet community. Biology, 10(2), 74.

Gatrell, A. C., Bailey, T. C., Diggle, P. J., and Rowlingson, B. S. (1996). Spatial point pattern analysis and its application in geographical epidemiology. Transactions of the Institute of British geographers, pages 256–274.

Giesen, C. et al. (2014). Highly multiplexed imaging of tumor tissues with subcellular resolution by mass cytometry. Nature methods, 11(4), 417–422.

Goldsmith, J., Crainiceanu, C. M., Caffo, B., and Reich, D. (2012). Longitudinal penalized functional regression for cognitive outcomes on neuronal tract measurements. Journal of the Royal Statistical Society: Series C (Applied Statistics), 61(3), 453–469.

Goltsev, Y. et al. (2018). Deep profiling of mouse splenic architecture with codex multiplexed imaging. Cell, 174(4), 968–981.

Heinzmann, K., Carter, L. M., Lewis, J. S., and Aboagye, E. O. (2017). Multiplexed imaging for diagnosis and therapy. Nature Biomedical Engineering, 1(9), 697–713.

Illian, J., Benson, E., Crawford, J., and Staines, H. (2006). Principal component analysis for spatial point processes—assessing the appropriateness of the approach in an ecological context. Case studies in spatial point process modeling, pages 135–150.

Ivanescu, A. E., Staicu, A.-M., Scheipl, F., and Greven, S. (2015). Penalized function-on-function regression. Computational Statistics, 30, 539–568.

Keren, L. et al. (2018). A structured tumor-immune microenvironment in triple negative breast cancer revealed by multiplexed ion beam imaging. Cell, 174(6), 1373–1387.

Kerscher, M. (2000). Statistical analysis of large-scale structure in the universe. In Statistical Physics and Spatial Statistics: The art of analyzing and modeling spatial structures and pattern formation, pages 36–71. Springer.

Legendre, P. and Fortin, M. J. (1989). Spatial pattern and ecological analysis. Vegetatio, 80, 107–138.

Lewis, S. M., Asselin-Labat, M.-L., Nguyen, Q., Berthelet, J., Tan, X., Wimmer, V. C., Merino, D., Rogers, K. L., and Naik, S. H. (2021). Spatial omics and multiplexed imaging to explore cancer biology. Nature methods, 18(9), 997–1012.

Lin, J.-R. et al. (2023). Multiplexed 3d atlas of state transitions and immune interaction in colorectal cancer. Cell, 186(2), 363–381.

Lin, J.-R., Fallahi-Sichani, M., and Sorger, P. K. (2015). Highly multiplexed imaging of single cells using a high-throughput cyclic immunofluorescence method. Nature communications, 6(1), 8390.

Liu, C. C. et al. (2022). Multiplexed ion beam imaging: insights into pathobiology. Annual Review of Pathology: Mechanisms of Disease, 17, 403–423.

Maoz, A., Dennis, M., and Greenson, J. K. (2019). The crohn’s-like lymphoid reaction to colorectal cancer-tertiary lymphoid structures with immunologic and potentially therapeutic relevance in colorectal cancer. Frontiers in Immunology, 10, 1884.

Nagtegaal, I. D. et al. (2020). The 2019 who classification of tumours of the digestive system. Histopathology, 76(2), 182.

Osher, N., Kang, J., Krishnan, S., Rao, A., and Baladandayuthapani, V. (2023). Spartin: a bayesian method for the quantification and characterization of cell type interactions in spatial pathology data. Frontiers in Genetics, 14, 1175603.

Ramsay, J. O. and Silverman, B. W. (2005). Functional Data Analysis. Springer.

Ripley, B. D. (1976). The second-order analysis of stationary point processes. Journal of applied probability, 13(2), 255–266.

Rozek, L. S. et al. (2016). Tumor-infiltrating lymphocytes, crohn’s-like lymphoid reaction, and survival from colorectal cancer. Journal of the National Cancer Institute, 108(8), djw027.

Schapiro, D. et al. (2017). histocat: analysis of cell phenotypes and interactions in multiplex image cytometry data. Nature methods, 14(9), 873–876.

Schürch, C. M. et al. (2020). Coordinated cellular neighborhoods orchestrate antitumoral immunity at the colorectal cancer invasive front. Cell, 182(5), 1341–1359.

Seal, S. and Ghosh, D. (2022). Miami: mutual information-based analysis of multiplex imaging data. Bioinformatics, 38(15), 3818–3826.

Seal, S., Wrobel, J., Johnson, A. M., Nemenoff, R. A., Schenk, E. L., Bitler, B. G., Jordan, K. R., and Ghosh, D. (2022). On clustering for cell phenotyping in multiplex immunohistochemistry (mihc) and multiplexed ion beam imaging (mibi) data. BMC Research Notes, 15(1), 215.

Shipkova, M. and Wieland, E. (2012). Surface markers of lymphocyte activation and markers of cell proliferation. Clinica Chimica Acta, 413(17-18), 1338–1349.

Smaga, L. and Zhang, J.-T. (2019). Linear hypothesis testing with functional data. Technometrics, 61(1), 99–110.

Tsakiroglou, A. M. et al. (2020). Spatial proximity between t and pd-l1 expressing cells as a prognostic biomarker for oropharyngeal squamous cell carcinoma. British Journal of Cancer, 122(4), 539–544.

Tsujikawa, T. et al. (2017). Quantitative multiplex immunohistochemistry reveals myeloid-inflamed tumor-immune complexity associated with poor prognosis. Cell reports, 19(1), 203–217.

Vipond, O. et al. (2021). Multiparameter persistent homology landscapes identify immune cell spatial patterns in tumors. Proceedings of the National Academy of Sciences, 118(41), e2102166118.

Vu, T., Wrobel, J., Bitler, B. G., Schenk, E. L., Jordan, K. R., and Ghosh, D. (2022). Spf: a spatial and functional data analytic approach to cell imaging data. PLOS Computational Biology, 18(6), e1009486.

Wallace, K. et al. (2021). Immune responses vary in preinvasive colorectal lesions by tumor location and histologyimmune responses in preinvasive lesions. Cancer Prevention Research, 14(9), 885–892.

Welch, W. J. (1990). Construction of permutation tests. Journal of the American Statistical Association, 85(411), 693–698.

Wilson, C. et al. (2022). Tumor immune cell clustering and its association with survival in african american women with ovarian cancer. PLoS Computational Biology, 18(3), e1009900.

Wrobel, J., Harris, C., and Vandekar, S. (2023). Statistical analysis of multiplex immunofluorescence and immunohistochemistry imaging data. In Statistical Genomics, pages 141–168. Springer.

Zhang, J.-T. (2013). Analysis of variance for functional data. CRC press.

