## Supplementary material for "SpaceANOVA: Spatial co-occurrence analysis of cell types in multiplex imaging data using point process and functional ANOVA" (from another version of this preprint: version 1, 2023-07-07)

### Additional results

In this supplementary material, we present a few additional figures and discussions pertinent to the analyses provided in the main text.

#### Real data analysis

##### Collection procedure for colorectal polyps dataset

The Medical University of South Carolina Hollings Cancer Center Biorepository and Tissue Analysis Shared Resource provided 5-micron thin sections for H&E and IF analysis. The study pathologist selected a convenience sample of 10 conventional colorectal adenomas (tubular, villous containing), 10 sessile serrated lesions, and 5 normal mucosal biopsies excised from patients undergoing colonoscopy between 2012-2014. All adenomas and serrated lesions were advanced size (1+cm). For all cases, the repository provided information on personal characteristics (age at diagnosis, sex, race) and clinicopathologic data (anatomic location, degree of dysplasia, histologic type) at diagnosis. For each case, using a newly created H&E, the study pathologist documented the histologic type (conventional adenoma (tubular (TA), villous (VA)) or sessile serrated adenoma (SSA)), degree of dysplasia, and other features of interest (e.g. lymphoid aggregate, necrosis). When there was disagreement in histologic type from the original diagnosis (n=4), the study pathologist's diagnosis was used to ensure uniformity.

##### Analysis of the type 1 diabetes dataset

Figures S1 and S2 correspond to the analysis of the T1DM dataset [3] described in the main manuscript.

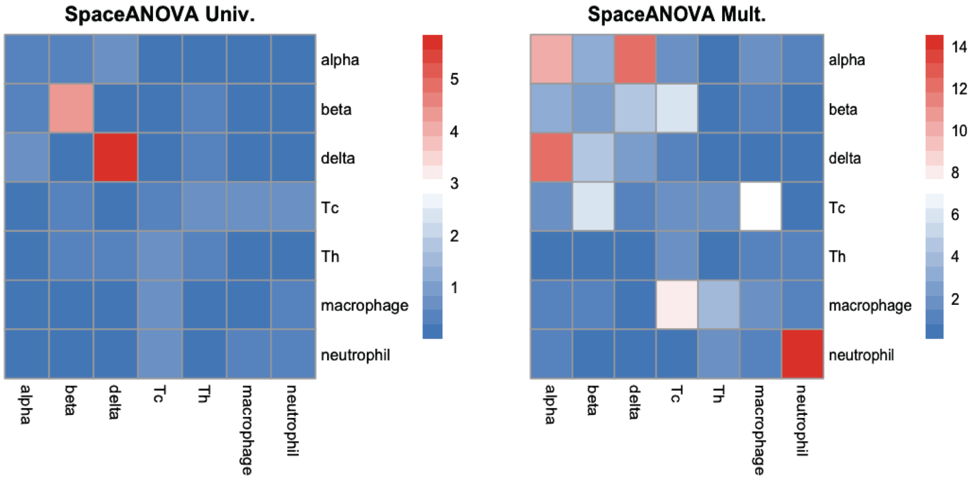


Figure S1: Heatmap of $-\log_{10}(p)$-values of the two methods for every pair of cell types.

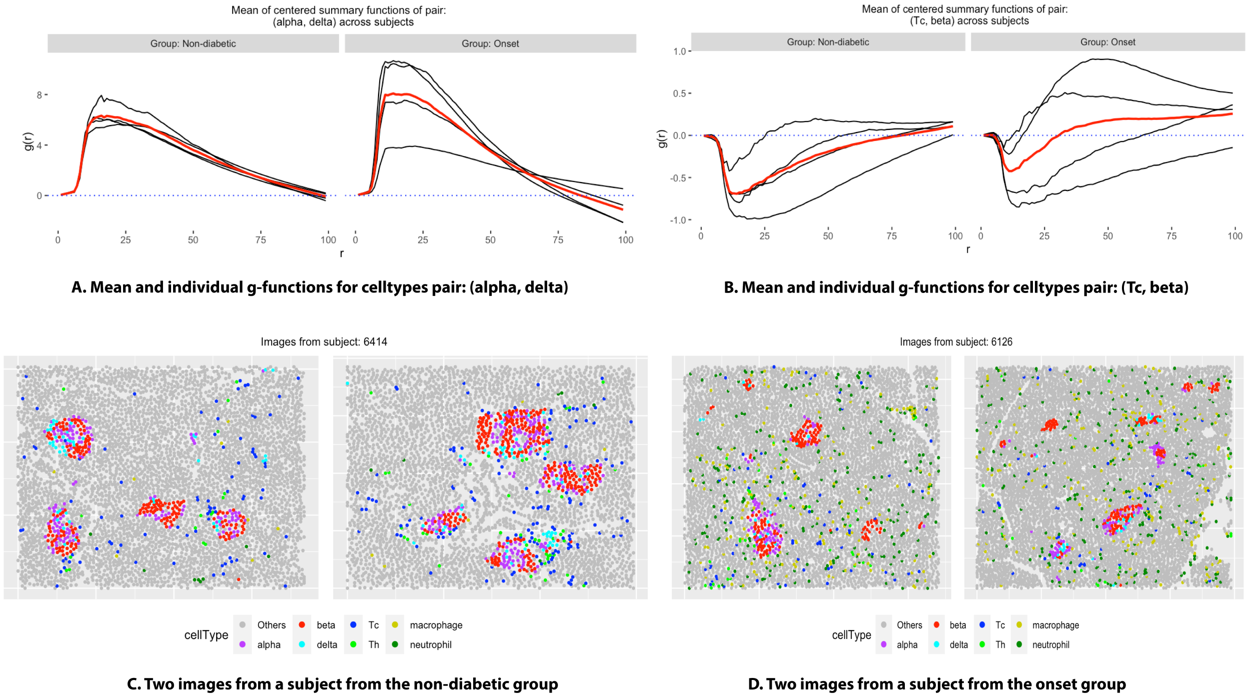


Figure S2: Figure (A) shows the individual g-function (averaged over images) of every subject in two groups (in the color black) and also the group mean function (in the color red), for the pair of cell types: (alpha, delta). Figure (B) shows a similar plot for the pair: (Tc, beta). Figures (C) and (D) show the cellular organization of representative images from two subjects from the groups non-diabetic and onset, respectively, where different colors correspond to different cell types.

#### Analysis of the CRC dataset

In the CRC dataset [4], there are two groups of subjects: one group (18 subjects) with the "Crohn's-like reaction" (CLR) tumor invasive front, while the other group (17 subjects) is defined by the presence of diffuse inflammatory infiltration (DII). Each subject has four images and cells of 29 different types of which we focused on studying the pairwise spatial co-occurrence of the top ten cell types in terms of cell count, namely, tumor-cells, cd68-cd163-macrophages, smooth-muscle, granulocytes, stroma, cd8˙t˙cells, cd4-t-cells-cd45ro, b-cells, vasculature, and cd4-t-cells. The range of $r$ was kept between [0, 100] with a step size of 10 which was higher than what we considered in the other two real data analyses because the number of cells was much lower in this dataset and a finer grid of $r$ would result in noisier and less smooth g-functions. For a similar reason and the tissue missing-ness not being a highlighted issue by Schutch et al. (2020), we considered unadjusted versions of SpaceANOVA Univ. and SpaceANOVA Mult. in this case.

Table S1: The pairs of cell types detected by SpaceANOVA Univ. for which the lowest $p$-values ($< 0.05$) were observed.

| cellType 1 | cellType 2 | p-value |
|---|---|---|
| tumor_cells | b_cells | 0.00017 |
| b_cells | tumor_cells | 0.00018 |
| tumor_cells | vasculature | 0.00050 |
| vasculature | tumor_cells | 0.00059 |
| cd68_cd163_macrophages | vasculature | 0.00689 |
| vasculature | cd8_t_cells | 0.00900 |
| vasculature | cd68_cd163_macrophages | 0.01051 |
| cd8_t_cells | vasculature | 0.01098 |
| cd68_cd163_macrophages | tumor_cells | 0.02287 |
| tumor_cells | cd68_cd163_macrophages | 0.02331 |
| smooth_muscle | tumor_cells | 0.02827 |
| tumor_cells | smooth_muscle | 0.03088 |
| tumor_cells | cd4_t_cells_cd45ro | 0.03280 |
| tumor_cells | cd8_t_cells | 0.03309 |
| cd8_t_cells | tumor_cells | 0.03340 |
| cd4_t_cells_cd45ro | tumor_cells | 0.03405 |
| granulocytes | granulocytes | 0.03820 |
| granulocytes | smooth_muscle | 0.03859 |
| cd4_t_cells_cd45ro | cd4_t_cells_cd45ro | 0.03915 |
| smooth_muscle | granulocytes | 0.04191 |
| stroma | tumor_cells | 0.04455 |
| tumor_cells | stroma | 0.04539 |
| cd8_t_cells | cd4_t_cells_cd45ro | 0.04741 |

In Tables S1 and S2, we display the lowest $p$-values and corresponding cell pairs as detected by

Table S2: The pairs of cell types detected by SpaceANOVA Mult. for which the lowest $p$-values ($< 0.05$) were observed.

| cellType 1 | cellType 2 | p-value |
|---|---|---|
| b_cells | granulocytes | 0.00550 |
| smooth_muscle | smooth_muscle | 0.00569 |
| granulocytes | b_cells | 0.00571 |
| cd68_cd163_macrophages | cd68_cd163_macrophages | 0.00817 |
| smooth_muscle | cd4_t_cells_cd45ro | 0.01747 |
| cd4_t_cells_cd45ro | smooth_muscle | 0.01986 |

SpaceANOVA Univ. and SpaceANOVA Mult., respectively. SpaceANOVA Univ. detected more pairs of cell types majority of which were also previously detected by SpicyR [2]. In Figure S3, we show the mean estimated g-functions corresponding to pairs: (b-cells, tumor-cells) and (vasculature, tumor-cells) which were detected by SpaceANOVA Univ., for all the subjects from the two groups: CLR and DII. We noted that for smaller values of $r$, the cell types from each of the pairs avoided each other in both CLR and DII. But, as $r$ increased the g-function became positive in the CLR group indicating a positive spatial co-occurrence, while in the DII group, the avoidance persisted. It can be partly explained by the fact that DII subjects have a very low count of b-cells. In the representative images from two subjects (Figure S3), one from each group, we noted that b-cells seemed to form prominent clusters in the CLR images, while tumor-cells and vasculatures were surrounded by themselves more in the DII images.

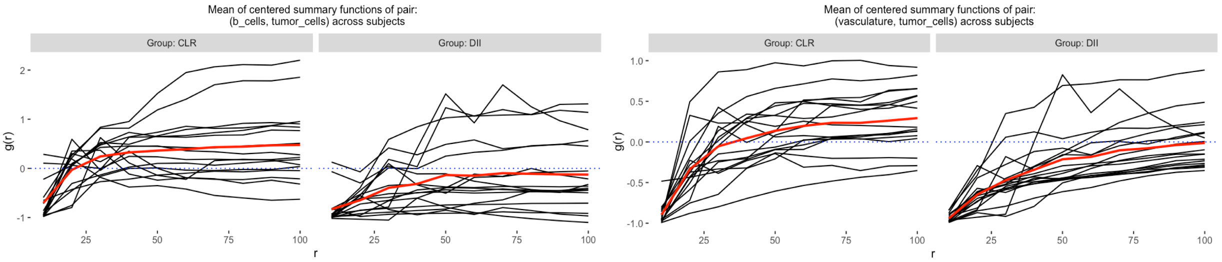


**A. Mean and individual g-functions for cell types pair: (b cells, tumor cells)**

**B. Mean and individual g-functions for cell types pair: (Vasculature, tumor cells)**

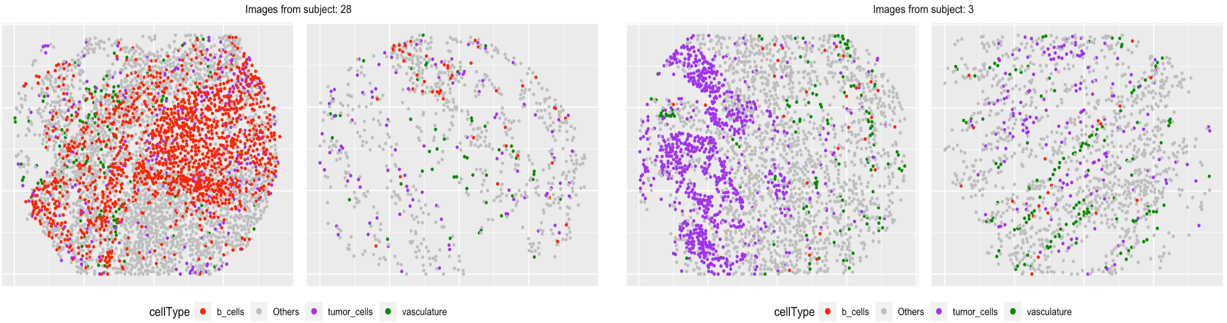


**C. Two images from a subject from the CLR group**

**D. Two images from a subject from the DII group**

Figure S3: Figure (A) shows the individual g-function (averaged over images) of every subject in two groups (in the color black) and also the group mean function (in the color red), for the pair of cell types: (b-cells, tumor-cells). Figure (B) shows a similar plot for the pair: (vasculature, tumor-cells). Figures (C) and (D) show the organization of four cell types in representative images from two subjects from the groups CLR and DII, respectively, where different colors correspond to different cell types.

#### Simulation plots

In Figure S4, we show how the cellular patterns look in two different simulation setups from Sections 4.1 and 4.2 of the main text. In Section 4.1, assumptions of a mixed Poisson point process previously outlined in [2] were used, while in Section 4.2, assumptions of a Neyman-Scott cluster process [1] were used. In Figure S4, the true $L$ and $g$ functions are displayed across varying values of the hyperparameters in the above two simulation setups.

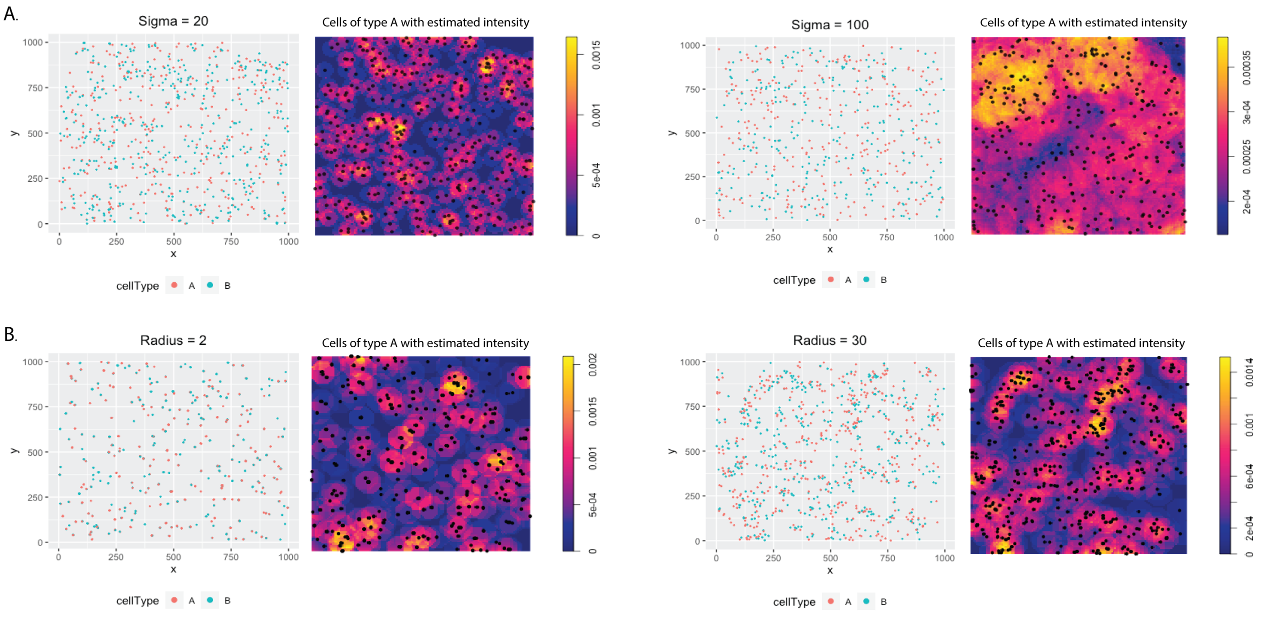


Figure S4: (A) corresponds to the simulation setup from Section 4.1 of the main text. It shows the organization of the two cell types along with the estimated density of the cell type A that was used to generate cells of type B. Notice that for a smaller value of the parameter Sigma, the co-occurrence is stronger (left panel). (B) shows a similar plot as (A) for the simulation setup from Section 4.2 of the main text. Notice that the co-occurrence is stronger for the larger value of the parameter Radius (right panel).

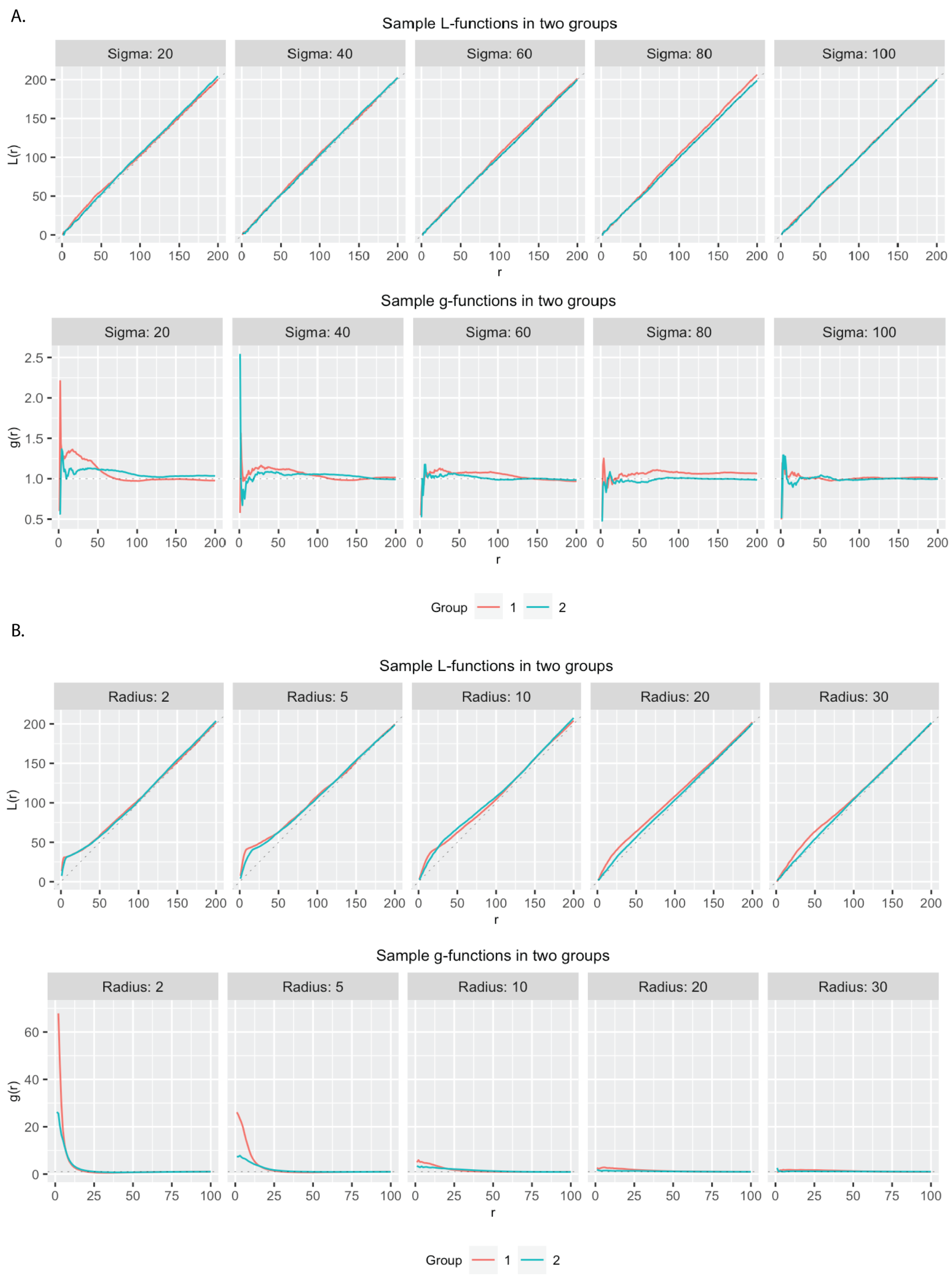


Figure S5: (A) shows the true L and g-functions for varying values of the parameter Sigma (and, Sigma-difference = Sigma) in the simulation setup from Section 4.1 of the main text. Notice that the difference between the functions of the two groups is decreasing as Sigma increases, more prominent for the L-functions. (B) shows a similar plot as (A) for varying values of the parameter Radius (and, Radius-difference = Radius) in the simulation setup from Section 4.2 of the main text. The difference between the functions of the two groups is becoming more apparent as Radius increases.

### Additional discussion about the method

#### The range of radius

In the first two real data analyses and simulation studies from the main text, we varied radius $r$ across an evenly spaced grid of values between [0, 100] and [0, 200], respectively, with step-size 1. In our experience, when the number of cells per image is not high enough, choosing a fine grid of $r$ could result in noisy summary functions. In such a case, the step size could be increased to a larger value such as 10 or 20 as we considered in the CRC data analysis which had only 258,385 cells in total as compared to 8,798,241, and 1,776,974 cells in the polyps and T1DM datasets, respectively. In addition, the association results may alter if different ranges of $r$ are considered. As an example, focusing only on the smaller values of $r$, say $r \in [0, 20]$ would reveal the occurrence of tight clusters. In general contexts, the range of $r$ is best chosen by pathological guidance and by exploring the data visually prior to fitting our model.

#### Distribution of the test statistics

Recall the definition of the two test statistics: $F_{univ}$ and $F_{mult}$ from the main text,

$$F_{univ} = \int_0^R \frac{\mathrm{SS}_G(r)/(G-1)}{\mathrm{SS}_E(r)/(N-G)} dr, \quad F_{mult} = \int_0^R \frac{\mathrm{SS}_G(r)/(G-1)}{\mathrm{SS}_E(r)/(N - \sum_{g=1}^G s_g)} dr.$$

Numerical approximations of these two statistics can be formulated as,

$$\widehat{F}_{univ} = \frac{1}{R_{len}} \sum_{i=1}^{R_{len}} \frac{\mathrm{SS}_G(r_i)/(G-1)}{\mathrm{SS}_E(r_i)/(N-G)}, \quad \widehat{F}_{mult} = \frac{1}{R_{len}} \sum_{i=1}^{R_{len}} \frac{\mathrm{SS}_G(r_i)/(G-1)}{\mathrm{SS}_E(r_i)/(N - \sum_{g=1}^G s_g)}$$

where $R_{len}$ denotes the number of grid values considered for radius $r$, and $r_i \in [0, R]$ denotes the $i$-th value in increasing order, with $r_1 = 0$ and $r_{R_{len}} = R$. To obtain the asymptotic null distribution of these two test statistics, we first need to estimate the covariance function $\gamma$. In the two cases, the pooled sample covariance functions can be defined as,

$$\widehat{\gamma}_{univ}(r_t, r_q) = \frac{1}{N-G}\sum_{g=1}^{G}\sum_{i=1}^{s_g}(Y_{gi}(r_t) - \bar{Y}_g(r_t))(Y_{gi}(r_q) - \bar{Y}_g(r_q))^T$$

$$\widehat{\gamma}_{mult}(r_t, r_q) = \frac{1}{N-\sum_{g=1}^{G} s_g}\sum_{g=1}^{G}\sum_{i=1}^{s_g}\sum_{j=1}^{n_{gi}}(Y_{gij}(r_t) - \bar{Y}_{gi}(r_t))(Y_{gij}(r_q) - \bar{Y}_{gi}(r_q))^T$$

Then, it can be shown that $\widehat{F}_{univ}$ and $\widehat{F}_{mult}$ approximately follow $\widehat{\beta}_1\chi^2_{\widehat{d}_1}$ and $\widehat{\beta}_2\chi^2_{\widehat{d}_2}$, respectively, where $\widehat{\beta}_1, \widehat{\beta}_2, \widehat{d}_1$, and $\widehat{d}_2$ have the following forms [5, 6],

$$\widehat{\beta}_1 = z_1\frac{tr(\widehat{\Gamma}^2_{univ})}{R},\ \widehat{d}_1 = \frac{1}{z_1^2}\frac{R^2}{tr(\widehat{\Gamma}^2_{univ})},\ \widehat{\Gamma}_{univ} = \left[\left[\frac{\widehat{\gamma}_{univ}(r_t, r_q)}{\sqrt{\widehat{\gamma}_{univ}(r_t, r_t)\widehat{\gamma}_{univ}(r_q, r_q)}}\right]\right], z_1 = \frac{N-G}{N-G-2}$$

$$\widehat{\beta}_2 = z_2\frac{tr(\widehat{\Gamma}^2_{mult})}{R},\ \widehat{d}_2 = \frac{1}{z_2^2}\frac{R^2}{tr(\widehat{\Gamma}^2_{mult})},\ \widehat{\Gamma}_{mult} = \left[\left[\frac{\widehat{\gamma}_{mult}(r_t, r_q)}{\sqrt{\widehat{\gamma}_{mult}(r_t, r_t)\widehat{\gamma}_{mult}(r_q, r_q)}}\right]\right], z_2 = \frac{N-\sum s_g}{N-\sum s_g - 2}.$$

#### The permutation adjustment

The permutation-based adjustment should be used if the images are known to have unnaturally missing holes. Using this adjustment may result in a loss of power, particularly if the number of cells is not large enough, but the results should be robust and indicative of only true associations.

#### Interpreting univariate vs. multivariate test results

Both SpaceANOVA Univ. and SpaceANOVA Mult. test the same hypothesis but treat the multiple g-functions (corresponding to multiple images) of a subject differently. Needless to mention that when every subject has exactly one image, these two methods collide. In all of our simulations, these methods performed very closely to each other with SpaceANOVA Mult. demonstrating slightly higher power. However, in a few cases, SpaceANOVA Mult. produced slightly inflated type-1 errors. It can be explained by the fact that the assumptions of SpaceANOVA Mult. are comparatively stringent since it does not acknowledge the higher-order dependencies between the g-functions of the same subject apart from sharing the same mean function. Thus, the unaccounted similarities might induce a small degree of bias in the results. SpaceANOVA Univ., on the other hand, does not suffer from this problem as it reduces the multiple functional profiles of a subject to a single profile and thus, does not require explicitly modeling the dependencies between those profiles.

It is worth understanding that SpaceANOVA Mult. operates on a larger sample size as it treats each individual image as a data point instead of just considering the subject-level mean data. This probably explains why it detected more pairs and the $p$-values were smaller compared to SpaceANOVA Univ. in the first two real data analyses. However, it should preferably be used in cases when the number of images per subject is large and more importantly, the images are "connected" or correspond to the same tissue or TME. This should be highlighted because in some scenarios, such as the CRC dataset [4], only a few specific highly disconnected regions of interest (just 4 in the case of the CRC dataset) are chosen from a subject's tissue. In such cases, the within-subject variation ($\mathrm{SS}_E(r)$ from Equation 4 of the main text) can become dominating, causing a loss of power for SpaceANOVA Mult. Intuitively, SpaceANOVA Mult. is somewhat similar to SpicyR-LM [2] which sheds the subject index entirely and focuses directly on the image-level data across groups. In SpicyR-Mixed, Canete et al. (2022) takes into account the subject index, ensuring that the images of a particular subject share some degree of dependency through random effect terms. These two highly different modeling approaches, however, result in entirely different conclusions, as seen in the SpicyR manuscript [2] and can thus be confusing to interpret.

### References

[1] Adrian Baddeley, Ege Rubak, and Rolf Turner. *Spatial point patterns: methodology and applications with R.* CRC press, 2015.

[2] Nicolas P Canete, Sourish S Iyengar, John T Ormerod, Heeva Baharlou, Andrew N Harman, and Ellis Patrick. spicyr: Spatial analysis of in situ cytometry data in r. *Bioinformatics*, 38(11):3099–3105, 2022.

[3] Nicolas Damond, Stefanie Engler, Vito RT Zanotelli, Denis Schapiro, Clive H Wasserfall, Irina Kusmartseva, Harry S Nick, Fabrizio Thorel, Pedro L Herrera, Mark A Atkinson, et al. A map of human type 1 diabetes progression by imaging mass cytometry. *Cell metabolism*, 29(3):755–768, 2019.

[4] Christian M Schürch et al. Coordinated cellular neighborhoods orchestrate antitumoral immunity at the colorectal cancer invasive front. *Cell*, 182(5):1341–1359, 2020.

[5] Łukasz Smaga and Jin-Ting Zhang. Linear hypothesis testing with functional data. *Technometrics*, 61(1):99–110, 2019.

[6] Jin-Ting Zhang and Xuehua Liang. One-way anova for functional data via globalizing the pointwise f-test. *Scandinavian Journal of Statistics*, 41(1):51–71, 2014.
